# PIKfyve/Fab1 is required for efficient V-ATPase and hydrolase delivery to phagosomes, phagosomal killing, and restriction of *Legionella* infection

**DOI:** 10.1101/343301

**Authors:** Catherine M. Buckley, Victoria L. Heath, Aurélie Guého, Cristina Bosmani, Paulina Knobloch, Phumzile Sikakana, Nicolas Personnic, Stephen K. Dove, Robert H. Michell, Roger Meier, Hubert Hilbi, Thierry Soldati, Robert H. Insall, Jason S. King

**Author notes:** Present address: Scientific Center for Optical and Electron Microscopy, ETH Zürich, CH-8093 Zürich, Switzerland.

## Abstract

By engulfing potentially harmful microbes, professional phagocytes are continually at risk from intracellular pathogens. To avoid becoming infected, the host must kill pathogens in the phagosome before they can escape or establish a survival niche. Here, we analyse the role of the phosphoinositide (PI) 5-kinase PIKfyve in phagosome maturation and killing, using the amoeba and model phagocyte *Dictyostelium discoideum*.

PIKfyve plays important but poorly understood roles in vesicular trafficking by catalysing formation of the lipids phosphatidylinositol (3,5)-bisphosphate (PI(3,5)_2_) and phosphatidylinositol-5-phosphate (PI(5)P). Here we show that its activity is essential during early phagosome maturation in *Dictyostelium*. Disruption of *PIKfyve* inhibited delivery of both the vacuolar V-ATPase and proteases, dramatically reducing the ability of cells to acidify newly formed phagosomes and digest their contents. Consequently, *PIKfyve*^-^ cells were unable to generate an effective antimicrobial environment and efficiently kill captured bacteria. Moreover, we demonstrate that cells lacking *PIKfyve* are more susceptible to infection by the intracellular pathogen *Legionella pneumophila*. We conclude that PIKfyve-catalysed phosphoinositide production plays a crucial and general role in ensuring early phagosomal maturation, protecting host cells from diverse pathogenic microbes.

**Importance:** Cells that capture or eat bacteria must swiftly kill them to prevent pathogens from surviving long enough to escape the bactericidal pathway and establish an infection. This is achieved by the rapid delivery of components that produce an antimicrobial environment in the phagosome, the compartment containing the captured microbe. This is essential both for the function of immune cells and for amoebae that feed on bacteria in their environment. Here we identify a central component of the pathway used by cells to deliver antimicrobial components to the phagosome and show that bacteria survive over three times as long within the host if this pathway is disabled. We show that this is of general importance for killing a wide range of pathogenic and non-pathogenic bacteria, and that it is physiologically important if cells are to avoid infection by the opportunistic human pathogen *Legionella*.

## Introduction

Professional phagocytes must kill their prey efficiently if they are to prevent the establishment of infections [1]. Multiple mechanisms are employed to achieve this. Once phagosomes have been internalised they quickly become acidified and acquire reactive oxygen species, antimicrobial peptides and acid hydrolases. The timely and regulated delivery of these components is vital to protect the host from intracellular pathogens but is incompletely understood.

After a particle is internalised, specific effector proteins are recruited to the phagosome's cytoplasmic surface by interacting with several inositol phospholipids (PIPs) that play important roles in regulating vesicle trafficking and controlling maturation. The effectors of each PIP regulate particular aspects of compartment identity, membrane trafficking and endosomal maturation [2, 3], Phosphatidylinositol-3-phosphate (PI(3)P), made by the class III PI 3-kinase VPS34, is one of the first PIPs to accumulate on vesicles after endocytosis, and P 1(3)P recruits proteins containing FYVE (Fab1, YOTB, Vac1 and EEA1) and PX domains, such as the canonical early endosome markers EEA1 and Hrs, and sorting nexins [4, 5]. Also recruited to early endosomes by its FYVE domain is PIKfyve (Fab1 in yeast) [6], a phosphoinositide 5-kinase that phosphorylates PI(3)P to phosphatidylinositol-3,5-bisphosphate (PI(3,5)P_2_) [7–11]. The roles of PI(3)P are well explored but the formation of PI(3,5)P_2_ and the identities and functions of its effector proteins and its metabolic product PI(5)P are less well understood [12–14]. PI(3,5)P_2_ is thought to accumulate predominantly on late endosomes, and disruption of PIKfyve activity leads to multiple endocytic defects, including gross endosomal enlargement and accumulation of autophagosomes [15–20]. Recent research has begun to reveal important physiological roles of PIKfyve and its products in a variety of eukaryotes, but mechanistic details remain sparse [21–25].

As in classical endocytosis, phagosome maturation is regulated by PIPs [26]. Phagosomes accumulate PI(3)P immediately after closure, and this is required for their subsequent maturation [27–29]. The recent identification of several PIKfyve inhibitors, including apilimod and YM201636 [30, 31], has allowed researchers to demonstrate the importance of PI (3) P to PI(3,5)P_2_ conversion for phagosomal maturation in macrophages and neutrophils [32–35]. However, there are conflicting reports on the roles of PIKfyve in key lysosomal functions such as acidification and degradation, with some studies reporting defective acidification upon PIKfyve inhibition [10, 36, 37] and others finding little effect [33, 38, 39]. Therefore the mechanistic roles of PIKfyve and its products, and their relevance to phagosome maturation, remain unclear and subject to debate.

To understand the function and physiological significance of PIKfyve, we have investigated its role in phagosome maturation and pathogen killing in *Dictyostelium discoideum*, a soil-dwelling amoeba and professional phagocyte that feeds on bacteria. *Dictyostelium* PIPs are unusual, with the lipid chain joined to the *sn*-1-position of the glycerol backbone by an ether, rather than ester, linkage: these PIPs should correctly be named as derivatives of plasmanylinositol rather than phosphatidylinositol [40]. This, however, makes no known difference to downstream functions, which are dictated by interactions between the inositol polyphosphate headgroups and effector proteins. *Dictyostelium* has thus been an effective model for analysis of phosphoinositide signalling [41–44]. For convenience, both the mammalian and *Dictyostelium* inositol phospholipids are referred to as PIPs hereafter.

We find that genetic or pharmacological disruption of PIKfyve activity in *Dictyostelium* leads to a swollen endosomal phenotype reminiscent of defects in macrophages. We provide a detailed analysis of phagosome maturation, and show that at least some of the defects in PIKfyve-deficient cells are due to reduced recruitment of the proton-pumping vacuolar (V-ATPase). Finally, we demonstrate that PIKfyve activity is required for the efficient killing of phagocytosed bacteria and for restricting the intracellular growth of the pathogen *Legionella pneumophila*.

## Results

### *PIKfyve Dictyostelium* have swollen endosomes

The *Dictyostelium* genome contains a single orthologue of *PIKfyve (PIP5K3)*. Like the mammalian and yeast proteins, *Dictyostelium* PIKfyve contains an N-terminal FYVE domain, a CCT (chaperonin Cpn60/TCP1)-like chaperone domain, a PIKfyve-unique cysteine/histidine-rich domain and a C-terminal PIP kinase domain [7]. In order to investigate the role of PI(3,5)P_2_ in *Dictyostelium* we disrupted the *PIKfyve* gene in the axenic Ax3 background by inserting a blasticidin resistance cassette and deleting a portion of the central PIKfyve-unique region (Supplementary Figure 1).

While the unusual ether-linked chemistry of the *Dictyostelium* inositol phospholipids prevented direct measurement of PI(3,5)P_2_ loss by either the standard method of methanolysis followed by HPLC of deacylation products or by mass spectrometry, we found that each mutant strain was highly vacuolated (Figure 1A and B), resembling the swollen vesicle phenotype observed upon *PIKfyve* knockdown or inhibition in mammalian cells, *C. elegans, S. cerevisiae* and *D. melanogaster* [10, 15, 20, 45]. This effect was recapitulated by incubation with the PIKfyve-specific inhibitor apilimod [30], confirming that this phenotype was due to deficient PIKfyve activity, most likely via the production of PI(3,5)P_2_ or PI(5)P (Figure 1C).

**Figure 1.**
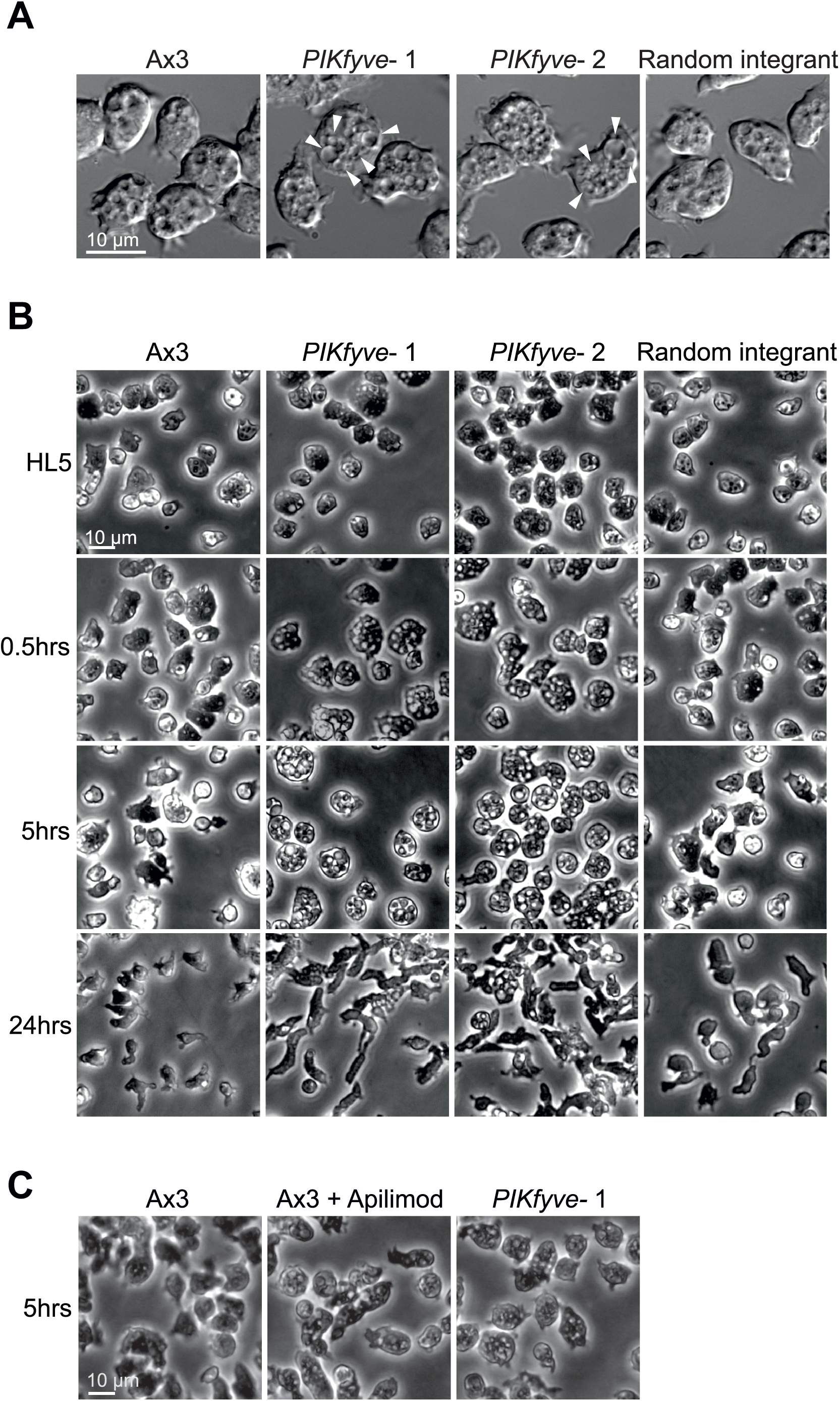
Knockout or inhibition of PIKfyve leads to a swollen vesicle phenotype. (A) DIC images of Ax3, two independent *PIKfyve^-^* clones and a random integrant growing in HL5 medium. Arrows indicate the enlarged vesicles. (B) Changes in morphology upon incubation in low osmolarity starvation buffer (KK2) compared to full medium (HL5). Swollen vesicles in *PIKfyve^-^* cells became initially more apparent but were lost as cells entered differentiation. (C) Induction of swollen vesicles with 5 μM apilimod, a PIKfyve-specific inhibitor, images taken in HL5 medium after 5 hours of treatment.

When *PIKfyve-* amoebae were hypotonically stressed in phosphate buffer, we observed a sustained increase in vacuolation for at least 5 hours. However, after 24 hours – when the cells became polarized indicating the onset of starvation-induced development – *PIKfyve-* mutants became indistinguishable from the random integrant and parental controls (Figure 1B). This is most likely due to the suppression of fluid-phase uptake that occurs when *Dictyostelium* cells enter starvation-induced development [46, 47]. Consistent with this interpretation, *PIKfyve*^-^ cells had no observable delay or other defects in development, and formed morphologically normal fruiting bodies with viable spores (Supplementary Figure 2). Therefore, disruption of PIKfyve leads to endocytic defects but it is not required for *Dictyostelium* development.

### PIKfyve is important for phagocytic growth but not uptake

Laboratory strains of *Dictyostelium* can obtain nutrients by macropinocytosis of liquid (axenic) medium or by phagocytosis of bacteria. Whilst *PIKfyve*^-^ cells had normal rates of both endocytosis and exocytosis, axenic growth was slower than for wild-type cells, with mutants doubling every 16 hours compared to 10 hours for the controls (Figure 2A-C).

**Figure 2.**
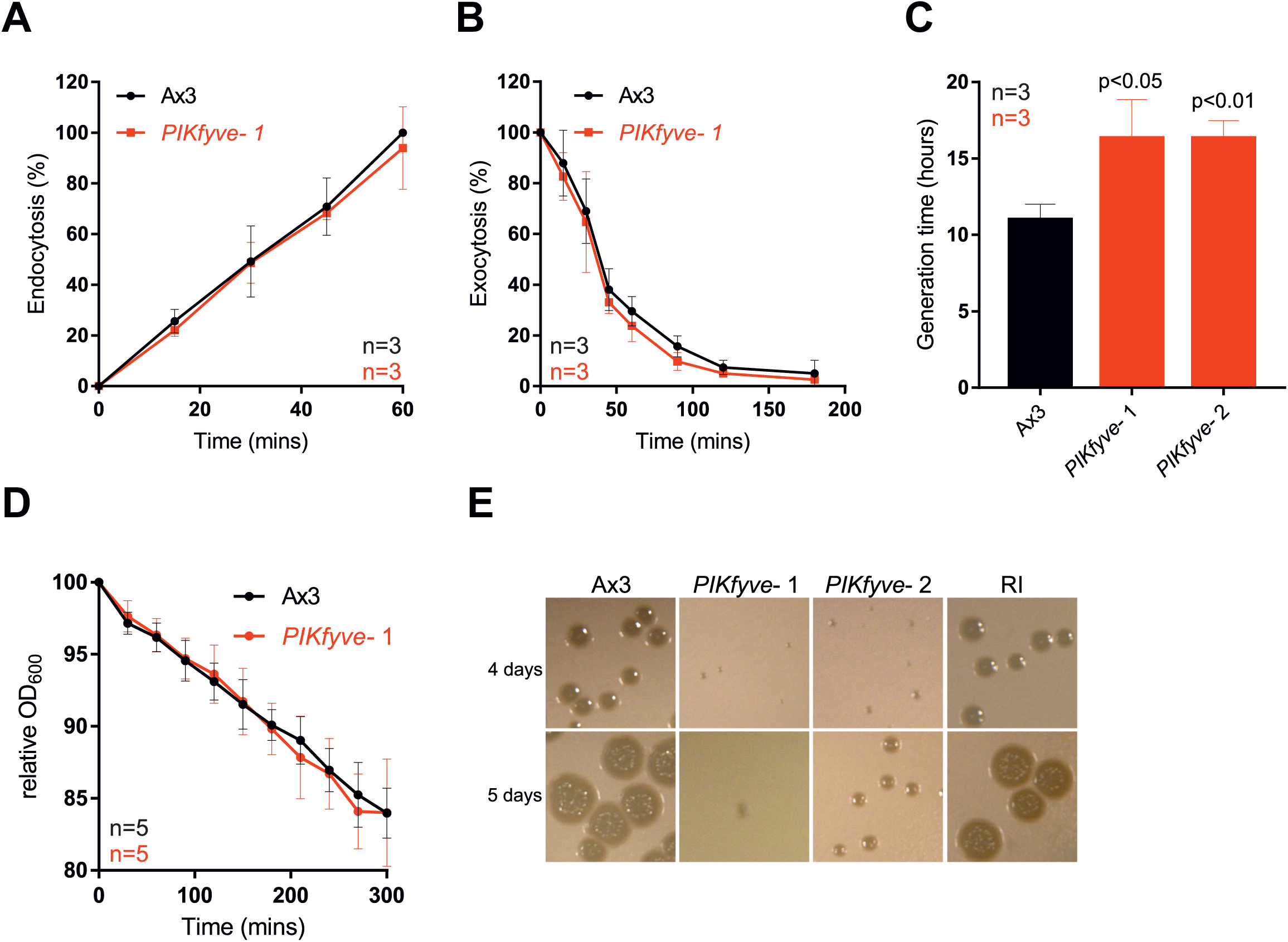
*PIKfyve*-null cells have growth defects. Measurement of (A) macropinocytosis or (B) exocytosis in Ax3 and *PIKfyve^-^* cells as measured by uptake or loss of FITC dextran. (C) Growth rates in axenic culture. *PIKfyve^-^* cells had a significantly longer generation time than Ax3 cells (Student's t-test). (D) Phagocytosis of *E. coli*, as measured by the ability of *Dictyostelium* cells to reduce the turbidity of a bacterial suspension. (E) Growth *PIKfyve* mutants on lawns of *K. pneumoniae* as indicated by the clearance of bacteria-free plaques on agar plates. All data plotted are mean +/-SD.

Growth on bacteria was more strongly affected. When we measured phagocytosis by following the ability of *Dictyostelium* cells to decrease the turbidity of an *E. coli* suspension over time we found no significant defects in *PIKfyve^-^* cells (Figure 2D). In contrast, they grew significantly more slowly on a lawn of *Klebsiella pneumoniae* (Figure 2E). This indicates a specific role for PIKfyve activity in phagosome maturation rather than bacterial uptake.

### *PIKfyve* deficient cells have defective phagosome acidification and V-ATPase delivery

Next we investigated how the absence of PIKfyve affects phagosomal maturation. One of the first stages of maturation is the acquisition of the proton-pumping V-ATPase, leading to rapid acidification [48, 49]. The influence of PIKfyve on endosomal pH regulation remains controversial: studies in *C. elegans*, plants, and mammalian epithelial cells have shown that PIKfyve is required for efficient acidification [37, 45, 50–52], but RAW 264.7 macrophages are still able to acidify their phagosomes to at least pH 5.5 when treated with a PIKfyve inhibitor [33].

Phagosome maturation is well characterised in *Dictyostelium*, with most studies being performed using mutants in the Ax2 genetic background, rather than Ax3 as used above. For comparison with other studies we isolated additional mutants from Ax2 cells, which were used for all subsequent experiments unless stated otherwise. Ax2 background *PIKfyve* mutants also exhibited slow growth on bacterial lawns but normal phagocytosis of both beads and bacteria (supplementary figure 3), confirming that these phenotypes are robust across multiple genetic backgrounds.

We followed phagosome acidification in *PIKfyve*^-^ *Dictyostelium* by measuring changes in the relative fluorescence of engulfed beads that had been labelled both with the pH-sensitive FITC and the pH-insensitive Alexa 594 succinimidyl ester [53]. Phagosomes from wild-type cells rapidly became acidified and remained acidic until ~40 minutes before transitioning to neutral post-lysosomes, but phagosomes of *PIKfyve*^-^ cells acidified much more slowly and never achieved as low a pH (Figure 3A).

**Figure 3.**
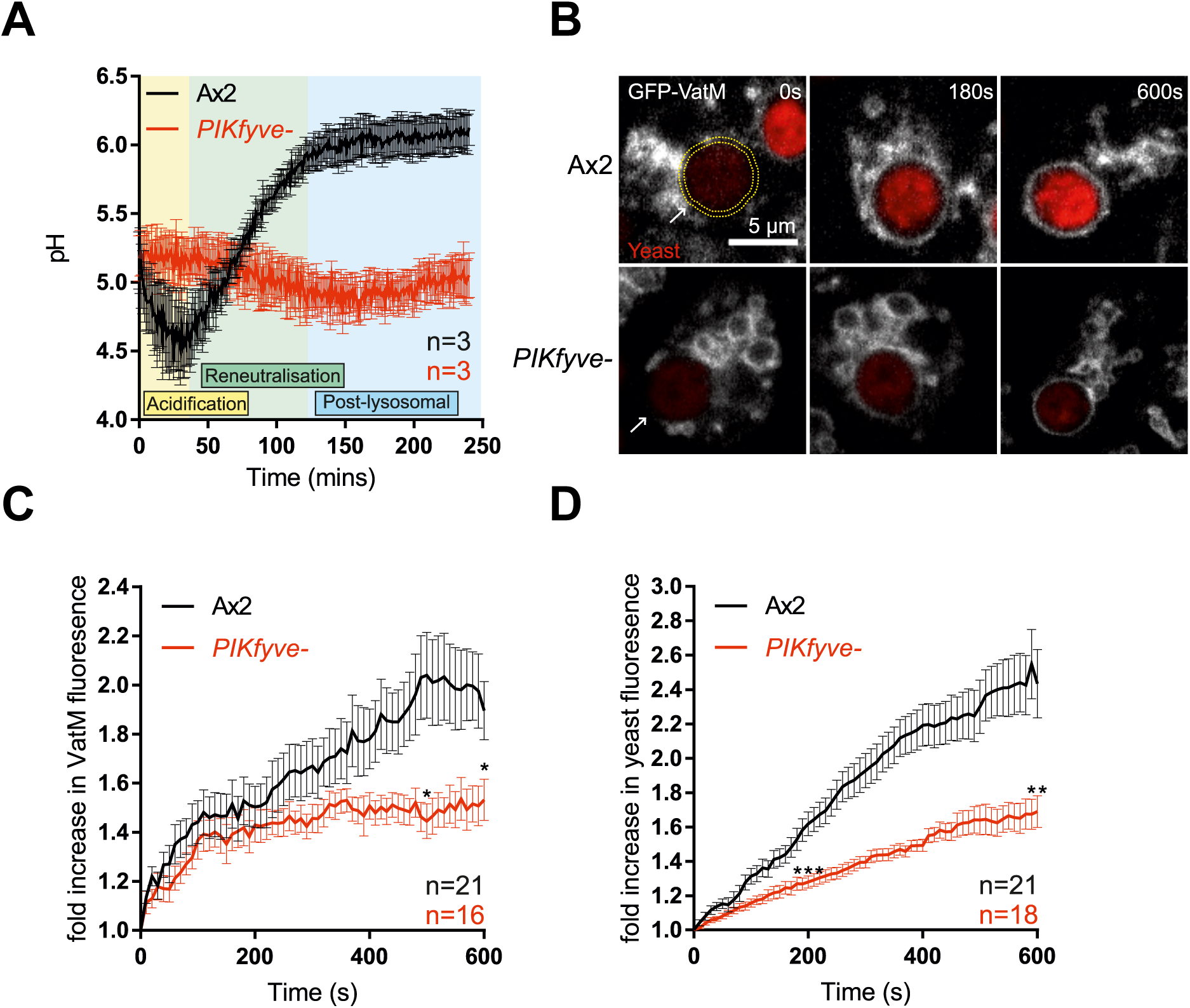
Disruption of *PIKfyve* prevents V-ATPase delivery and phagosome acidification. (A) Measurement of phagosomal acidification in Ax2 and *PIKfyve^-^* cells after engulfment of beads conjugated to pH-sensitive fluorophores. After initial acidification, *Dictyostelium* phagosomes reneutralise ~45 minutes prior to exocytosis. (B) Recruitment of the V-ATPase subunit GFP-VatM to phagocytosed pHrodo-labelled yeast visualised by confocal time-lapse imaging. (C) Quantification of GFP-VatM recruitment over time averaged across multiple phagocytic events. Images were quantified by automated selection of a 0.5 μm-thick ring surrounding the yeast at each time point (see yellow dotted circle in (B)). N indicates the total number of cells analysed over 3 independent experiments. Quantification of the associated increase in yeast-associated pHrodo fluorescence, indicating acidification is shown in (D). Data shown are mean +/-SEM, p values calculated by Student's t-test.

Phagosomal acidification is driven by the rapid recruitment and activity of the V-ATPase. To differentiate between defective V-ATPase delivery and activity, we directly imaged recruitment of the VatM transmembrane subunit of the V-ATPase fused to GFP to nascent phagosomes by microscopy. By observing the phagocytosis of pH-sensitive pHrodo-labelled yeast we were able to simultaneously monitor acidification.

GFP-VatM began accumulating on phagosomes immediately following internalisation both in *PIKfyve*^-^ and control cells, but it accumulated more slowly in the mutants and to only about half of the levels observed in wild-type cells (Figure 3B and C). Defective acidification was further demonstrated by a reduced increase in pHrodo fluorescence (Figure 3D). We conclude that there is some PIKfyve-independent lysosomal fusion with phagosomes, but that PIKfyve activity is required for the high levels of V-ATPase accumulation that are needed for efficient phagosomal acidification.

The V-ATPase consists of V_0_ (transmembrane) and V_1_ (peripheral) subcomplexes. It has previously been suggested that PI(3,5)P_2_ can regulate V_0_-V_1_ assembly at the yeast vacuole allowing dynamic regulation of activity [54]. VatM is a component of the V_0_ subcomplex (subunit a in mammals and yeast). To test whether V_0_-V_1_ association is also affected by loss of PIKfyve we observed the phagosomal recruitment of the V_1_ subunit VatB fused to GFP [55]. Both GFP-VatM and VatB-GFP were expressed equally in wild-type and mutant cells (Supplementary Figure 4A). As before VatB-GFP recruitment to nascent phagosomes was also significantly decreased in *PIKfyve-* cells (Supplementary Figure 4B and C)

**Figure 4.**
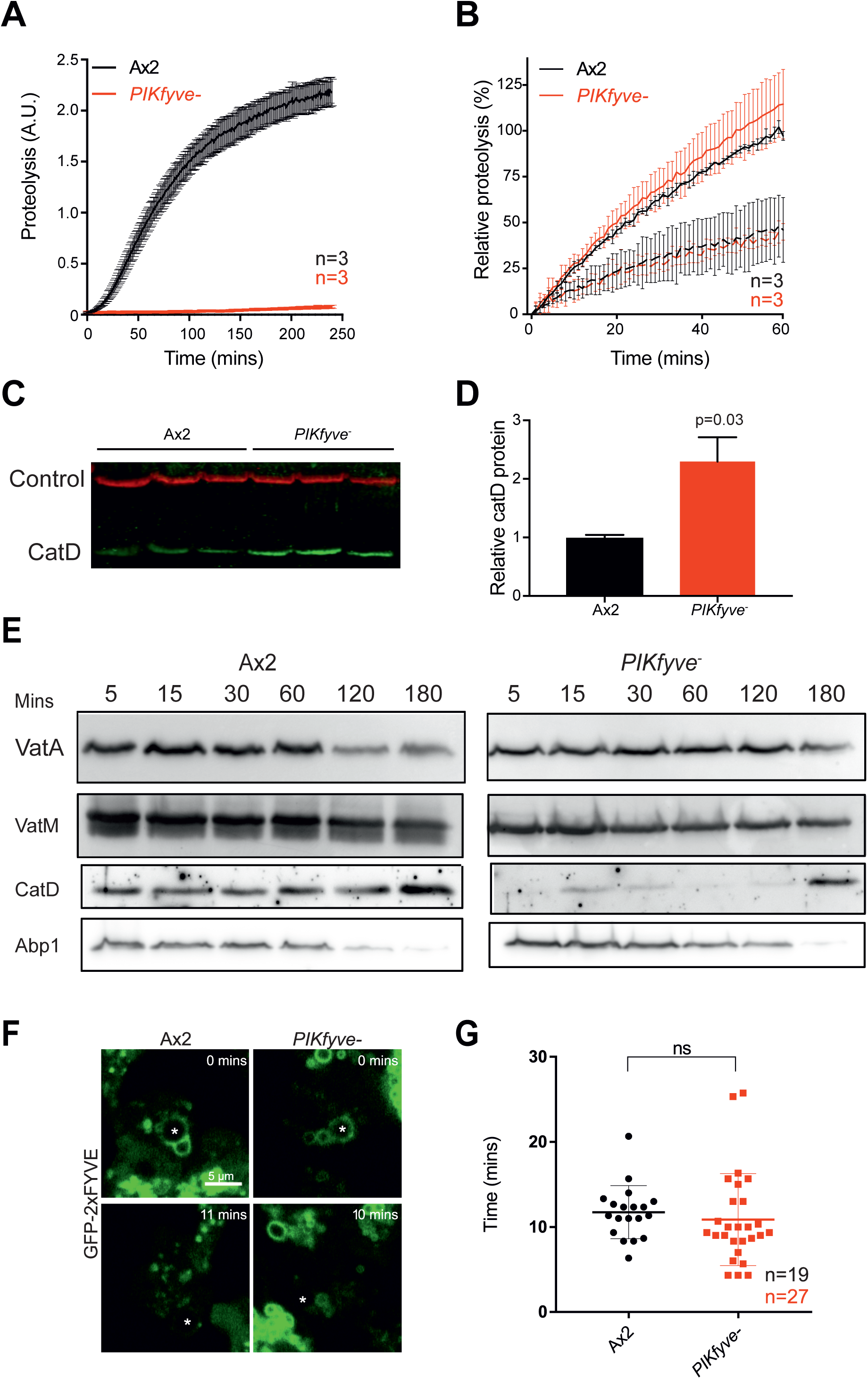
PIKfyve is required for hydrolase activity and proteolysis. (A) Phagosomal proteolysis measured by dequenching of DQ-BSA-conjugated beads after phagocytosis. (B) Total proteolytic activity in cell lysates against DQ-BSA-beads is unchanged upon *PIKfyve* disruption, dotted lines are parallel samples in the presence of protease inhibitors. (C) Western blot of cathepsin D protein levels, three independent samples of each strain were normalised for total protein and are quantified in (D). Loading control is fluorescent streptavidin which recognises the mitochondrial protein MCCC1; P-value from a one-sample T-test. (E) Analysis of phagosome maturation, by purifying phagosomes from cells after maturation for the indicated times. VatA is a subunit of the Visubcomplex of the V-ATPase, whereas VatM is a V_0_ component. Blots are from the same samples and are representative of multiple independent purifications. (F) Analysis of PI(3)P dynamics in the absence of PIKfyve. PI(3)P was monitored by the recruitment of GFP-2xFYVE following phagocytosis of 3 μm beads (asterisks) imaged by confocal time-lapse microscopy. (G) Time that GFP-2xFYVE stays associated with phagosomes following engulfment, indicating that PI(3)P removal is not PIKfyve-dependent. N indicates the total number of cells quantified across 3 independent experiments. Data shown are mean ± SEM (A & B) or SD (G).

It should be noted that expression of VatB-GFP (but not GFP-VatM) caused a partial inhibition of acidification in this assay, indicating caution should be taken in using this construct (Supplementary Figure 4D). Nevertheless, the observation that both V-ATPase components were equally affected by PIKfyve deletion suggests that PIKfyve is required for delivery of the entire V-ATPase to the phagosome, rather than specifically affecting V_0_-V_1_ association.

### Proteolytic activity and hydrolase delivery are specifically affected in *PIKfyve*^-^ cells

Proper degradation of internalised material requires both acidification and the activity of proteases. To test if hydrolytic activity was also dependent on PIKfyve, we measured phagosomal proteolysis by following the increase in fluorescence due to the cleavage and unquenching of DQ-bovine serum albumin (DQ-BSA) coupled to beads [53] (Figure 4A). Strikingly, despite their partial acidification, phagosomes in *PIKfyve* cells exhibited an almost complete loss of proteolytic activity. To control for a potential general decrease in protease activity, we measured the unquenching of DQ-BSA beads in whole cell lysates (Figure 4B) and the protein levels of lysosomal protease cathepsin D by Western blot (Figure 4C and D). Although activity in the phagosome is completely lost, we found that total lysosomal activity remained normal and cathepsin D levels were significantly increased upon loss of *PIKfyve*, consistent with a defect in delivery to the phagosome, rather than in lysosomal biogenesis.

To investigate whether PIKfyve activity was required to deliver proteases to the phagosome, we purified phagosomes at different stages of maturation and analysed their composition by Western blot (Figure 4E). In wild-type cells phagosomes acquired cathepsin D from the earliest time-point, but the protease was almost completely absent from phagosomes in *PIKfyve^-^* cells. Whilst the delivery of the vacuolar ATPase was also qualitatively reduced in this assay, consistent with decreased acidification, Actin-binding protein 1 (Abp1), an independent marker of phagosome maturation [56], was unaffected. Thus, although both hydrolase and to a lesser extent V-ATPase delivery requires PIKfyve activity, other aspects of maturation appear to progress normally.

### PIKfyve does not regulate PI(3)P dynamics

We next wanted to confirm whether loss of PIKfyve disrupts specific aspects of phagosome maturation or causes a general trafficking defect. PI(3)P is one of the best characterized early markers of maturing endosomes and phagosomes in both mammalian macrophages and *Dictyostelium*. Immediately following particle internalization, PI(3)P is generated on the phagosome by the class III PI 3-kinase VPS34 [27, 33, 57] and interacts with a number of important regulators of maturation such as Rab5 [58]. PIKfyve is both recruited by PI(3)P and phosphorylates it, forming PI(3,5)P_2_. Loss of PIKfyve activity could perturb phagosome maturation by reducing PI(3)P consumption, by eliminating the actions of PI(3,5)P_2_, or both. Studies on macrophages have indicated that inhibition of PIKfyve can cause prolonged PI(3)P signalling [33].

PI(3)P can be visualized in cells using the well-characterised reporter GFP-2xFYVE [41, 59]. Expression of GFP-2xFYVE in control cells demonstrated that PI(3)P is present on *Dictyostelium* phagosomes immediately following engulfment, consistent with previous reports [57] (Figure 4F). However, we found no abnormalities in either the recruitment to or the dissociation from phagosomes of this reporter in *PIKfyve^-^* mutants (Figure 4F and G). PIKfyve activity seems not to influence the steady-state levels of PI(3)P in *Dictyostelium*, and the functional defects in *PIKfyve^-^* cells therefore are likely to be caused by a lack of the product(s) of PIKfyve activity.

Overall, the normal transition of *PIKfyve^-^* phagosomes into a PI(3)P-negative compartment indicates that much of their phagosome maturation process continues normally - but with the product(s) of PIKfyve action playing specific role(s) in the delivery of certain important components, including the V-ATPase and hydrolytic enzymes.

### PIKfyve is essential for efficient killing of bacteria

Acidification and proteolysis are important mechanisms used by phagocytes to kill engulfed microbes. We therefore asked whether PIKfyve was physiologically important for killing. Bacterial death leads to membrane permeabilisation and intracellular acidification, so survival time within phagosomes can be inferred by observing the phagocytosis and subsequent quenching of GFP fluorescence when expressed by a non-pathogenic *Klebsiella pneumoniae* strain [60]. In this assay, the phagocytosed bacteria survived more than three times longer in *PIKfyve*^-^ cells (median survival 12 min) than in wild-type cells (3.5 min) (Figure 5A and B). These benign bacteria did eventually die in *PIKfyve*^-^ cells, indicating either that the residual acidification is eventually sufficient or that enough other elements of the complex bacterial killing machinery remain functional in *PIKfyve*^-^ phagosomes. These defects in bacterial killing and digestion (Figure 4) explain why PIKfyve is important for *Dictyostelium* to grow on a lawn of *K. pneumoniae* (Figure 2E). To test whether this is general to a broad range of bacteria, we employed an assay whereby serial dilutions of amoebae are plated on lawns of a panel including both Gram-positive and Gram-negative bacteria [61]. In this assay, *PIKfyve*-deficient cells were severely inhibited in growth on all bacteria tested, demonstrating a general role for PIKfyve in bacterial killing and digestion.

**Figure 5.**
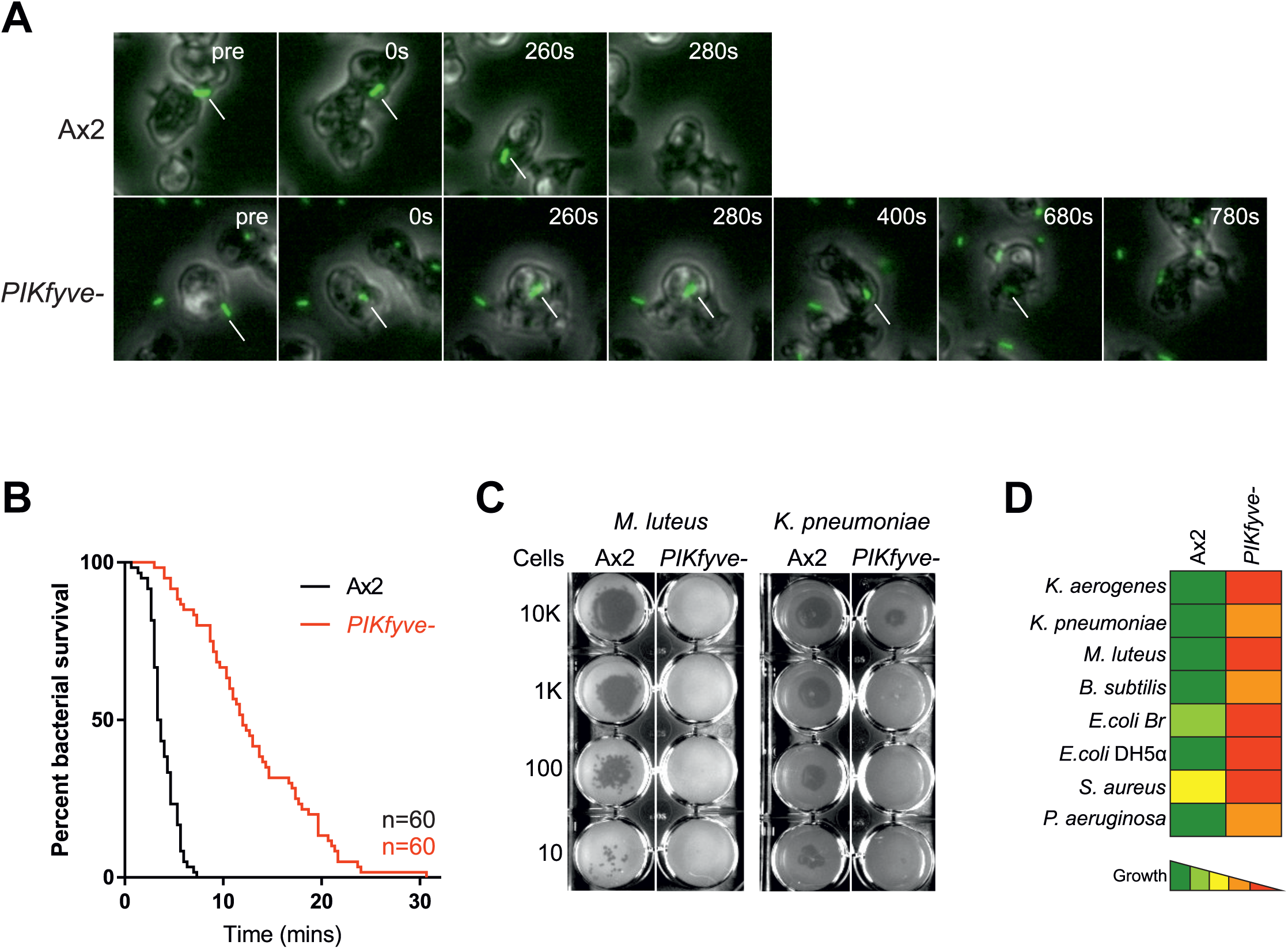
Bacterial survival is increased in *PIKfyve*-null cells. (A) Stills from widefield timelapse movies of *Dictyostelium* cells eating GFP-expressing *Klebsiella pneumoniae*. The point of bacterial cell permeablisation and death can be inferred from the quenching of GFP fluorescence after engulfment. Arrows indicate captured bacteria. This is quantified in (B) which shows a Kaplan-Meyer survival graph, based on the persistence of bacteria GFP-fluorescence within the amoebae. 60 bacteria were followed across three independent experiments and survived significantly longer in *PIKfyve^-^*null cells than Ax2 (p<0.0001, Mantel-Cox test). (C) Loss of PIKfyve inhibits growth on diverse bacteria. Growth was assessed by plating serial dilutions of amoebae on lawns of different bacteria and dark plaques indicate amoebae growth. Data for all bacteria are summarised in (D).

### PIKfyve activity restricts the persistence of *Legionella* infection

Many pathogenic bacteria infect host immune cells by manipulating phagosome maturation to establish a replication-permissive niche or to escape into the cytosol. To avoid infection, host cells must efficiently kill such pathogens; hence PIKfyve might be critical to protect host cells from infection.

*Legionella pneumophila* is a Gram-negative opportunistic human pathogen that normally lives in the environment by establishing replicative niches inside amoebae such as *Acanthamoeba*. This process can be replicated in the laboratory using *Dictyostelium* as an experimental host [62]. Following its phagocytosis, *Legionella* can disrupt normal phagosomal maturation and form a unique *Legionella-* containing vacuole (LCV). This requires the Icm/Dot (Intracellular multiplication/Defective for organelle trafficking) type IV secretion system that delivers a large number of bacterial effector proteins into the host (reviewed in [63]). These effectors modify many host signalling and trafficking pathways, one of which prevents the nascent *Legionella-*containing phagosome from fusing with lysosomes [64].

Phosphoinositide signalling is heavily implicated in *Legionella* pathogenesis, with *Legionella-* containing phagosomes rapidly accumulating PI(3)P. Its concentration then declines within 2 hours and PI(4)P accumulates [65]. Multiple effectors introduced through the Icm/Dot system bind PI(3)P or PI(4)P [66–72]. The role of PI(3,5)P_2_ (and/or PI(5)P) in *Legionella* infection is yet to be investigated, so we tested whether PIKfyve was beneficial or detrimental for the host to restrain *Legionella* infection.

When we measured the rate of uptake of GFP-expressing *Legionella* by flow cytometry, we found *PIKfyve*^-^ *Dictyostelium* were indistinguishable from wild-type. Both strains phagocytosed many more of the virulent wild-type *Legionella* strain (JR32) than an avirulent strain that is defective in type IV secretion (∆*icmT*) [73] (Figure 6A). These results are in agreement with the previous finding that expression of the Icm/Dot T4SS promotes uptake of *Legionella* [74]. When we measured the ability of *Dictyostelium* to kill *Legionella* ∆*icmT*, the bacteria survived for significantly longer in *PIKfyve^-^* cells (Figure 6B).

**Figure 6.**
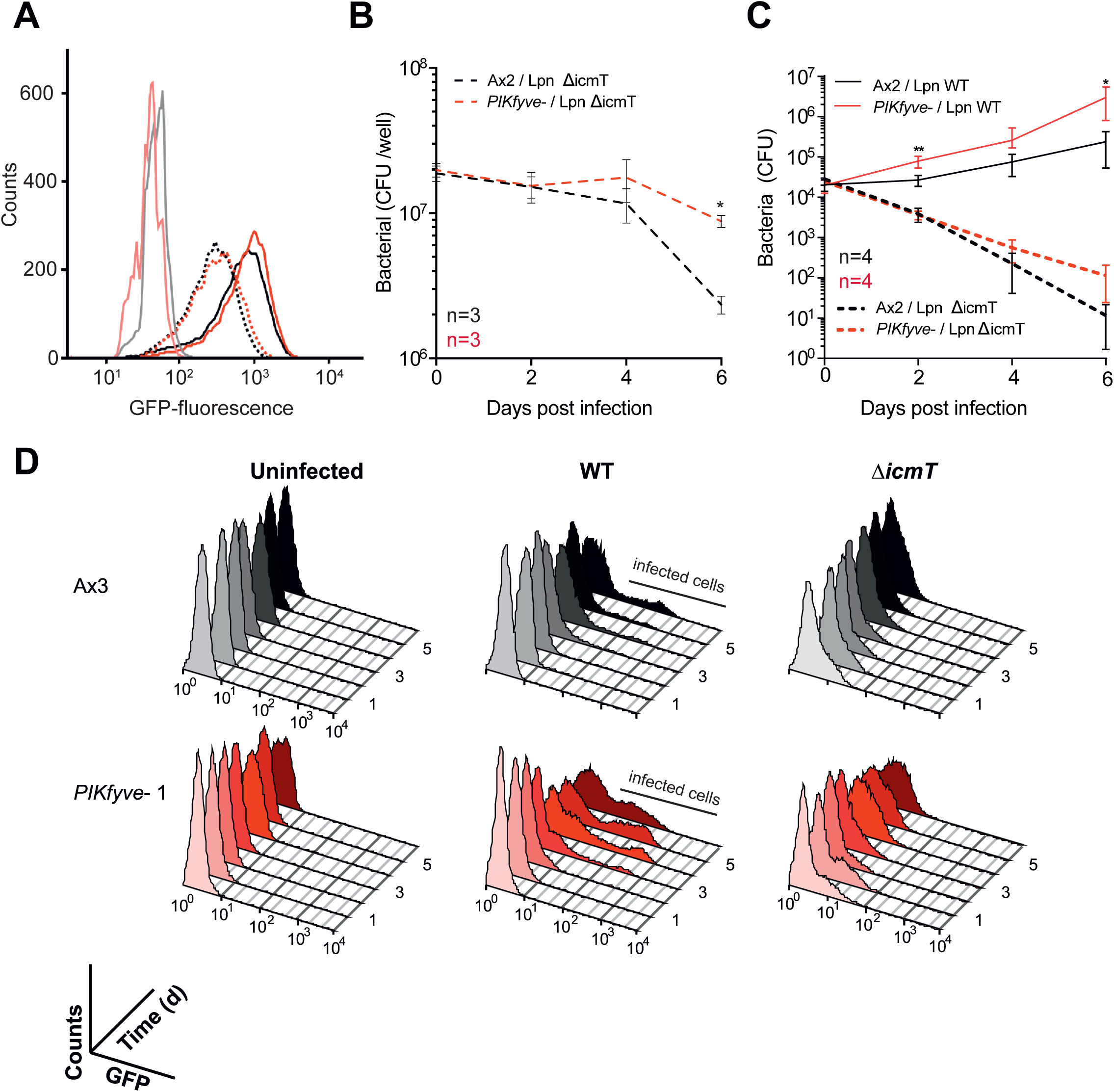
PIKfyve is required to suppress *Legionella* replication. A) Wild-type (Ax2, black lines), or *PIKfyve^-^*null *Dictyostelium* (red lines) were infected (MOI 50) with wild-type (JR32, solid lines) or avirulent (∆*icmT*, dotted lines) *Legionella* expressing GFP and fixed 40 min post infection before analysis by flow cytometry. The GFP fluorescence intensity, indicating bacteria per cell was indistinguishable between the two *Dictyostelium* strains, but higher for JR32 than ∆*icmT*. Uninfected cells are represented by pale black/red lines. Data are representative of three independent experiments, performed in duplicate. (B) Survival of GFP*-*∆*icmT Legionella* after infecting Ax2 and *PIKfyve^-^*null amoebae (MOI 50). Bacterial colony forming units (CFU) were determined at each timepoint after lysis of the *Dictyostelium* amoebae. (C) Outcome of infection with either wild-type (solid lines) or ∆*icmT* (dashed lines) *Legionella. Dictyostelium* cells were infected at a MOI of 0.1, and intracellular growth measured by CFU's at each indicated time. Data shown are the means +/-SEM of 3 independent experiments performed in triplicate (*=p<0.05, **p<0.01 Student's t-test vs equivalent Ax2 infection). (D) Flow cytometry of intracellular bacterial burden of Ax3-derived *PIKfyve^-^*null cells infected with GFP-producing *Legionella* strains over time. Virulent *Legionella* replicate more efficiently in *PIKfyve*^-^ *Dictyostelium*, as indicated by the increasing proportions of amoebae containing high levels of GFP over time. Graphs show >10,000 cells measured at each time point, and are representative of 3 independent experiments.

We next tested the role of PIKfyve on the outcome of infection. When Ax2 and PIKfyve mutants were infected with either wild-type or *∆icmT Legionella*, using a MOI of 0.1 to compensate for the greater uptake of wild-type *Legionella*, both amoeba strains suppressed the avirulent bacteria, although the reduction in bacteria was slower in the *PIKfyve* mutants. In contrast, wild-type *Legionella* grew substantially faster in cells lacking PIKfyve (Figure 6C, note that the CFUs scales in Figure 6B and C are logarithmic). We independently confirmed these results using flow cytometry of cells infected with GFP-producing bacteria in our Ax3-background mutants. The only *Dictyostelium* cells that accumulated substantial GFP fluorescence over several days were those infected by wild-type *Legionella*, and this happened sooner and to a greater degree in the *PIKfyve^-^* cells (Figure 6D).

Unlike PI(3)P and PI(4)P, the lipid products of PIKfyve are thus not required for *Legionella* to subvert phagosome maturation and generate its replicative vacuole. Rather, the role of PIKfyve in ensuring rapid phagosomal acidification and digestion is crucial for the host to prevent *Legionella*, and presumably other pathogens, from surviving and establishing a permissive niche.

## Discussion

In this study, we have characterised the role of PIKfyve during phagosome maturation using the model phagocyte *Dictyostelium*. The roles of PI 3-kinases and PI(3)P signalling during phagosome formation and early maturation have been studied extensively but the subsequent actions of PIKfyve and roles of PI(3,5)P_2_ and PI(5)P are much less well understood [3, 26]. In non-phagocytic cells such as fibroblasts and yeast, PI(3,5)P_2_ production is important for endosomal fission and fragmentation of endolysosomal compartments [10, 18, 37, 45], and PIKfyve inhibition has been shown to cause macrophage lysosomes to coalesce by an unknown mechanism [34]. PIKfyve also regulates macropinosome maturation [38], intracellular replication of the *Vaccinia* virus and *Salmonella* [38, 51, 75] and production of reactive oxygen species (ROS) in neutrophils [35]. In this paper we show that PIKfyve is critical to ensure efficient phagosomal acidification and proteolysis via delivery of specific components, and we demonstrate its physiological importance in the killing of bacteria and suppression of intracellular pathogens.

Complex effects of PIKfyve inhibition on PIP-mediated signalling have hampered clear interpretation of PIKfyve function in some mammalian studies. For example, some studies report that disruption of PIKfyve both prolonged PI(3)P-mediated signalling and eliminated PI(3,5)P_2_ production [30, 32], making it difficult to determine which phosphoinositide change is responsible for the observed phenotypes. In contrast, and in agreement with other reports in mammalian cells [18, 37], we found that deletion of PIKfyve had no impact on phagosomal PI(3)P dynamics in *Dictyostelium*. The defects in phagosome maturation that we observed in this system are thus due to lack of PI(3,5)P_2_/PI(5)P formation and not to prolonged PI(3)P signalling. There is limited evidence that PIKfyve might exhibit protein kinase activity [76], but whether this is relevant *in vivo* remains to be shown.

The role of PIKfyve in lysosomal acidification and degradation is currently disputed. Several studies which have measured vesicular pH at a single time point have shown that PIKfyve is required for acidification [10, 37, 45, 52], whereas others found that disruption of PIKfyve had little effect on phagosomal pH [33, 38, 39]. In contrast, we followed the temporal dynamics of V-ATPase delivery and of phagosomal acidification and proteolysis, and showed that V-ATPase delivery to PIKfyve-deficient phagosomes was substantially decreased and delayed, with consequent defects on initial acidification and proteolysis. PI(3,5)P_2_ has also been proposed to regulate V-ATPase V_0_-V_1_ subcomplex association dynamically at the yeast vacuole [54], but we found no evidence for this during *Dictyostelium* phagosome maturation.

It is still not clear how PIKfyve-generated PI(3,5)P_2_ regulates V-ATPase trafficking, and few PI(3,5)P_2_ effectors are known. One of these is the lysosomal cation channel TRPML1/mucolipin, which is specifically activated by PI(3,5)P_2_ [77]. This interaction was recently shown to partly underlie the role of PIKfyve in macropinosome fragmentation, although not acidification [38]. TRPML1 is also required for phagosome-lysosome fusion [78], and PI(3,5)P_2_ and TRPML1 have been proposed to mediate interactions between lysosomes and microtubules [79]. PIKfyve may therefore drive V-ATPase delivery to phagosomes both by microtubule-directed trafficking and by regulating fission. However, the sole mucolipin orthologue in *Dictyostelium* is only recruited to phagosomes much later, during the post-lysosomal phase, and its disruption influences exocytosis rather than acidification [80].

Effective phagosomal acidification and proteolysis is essential if phagocytes are to kill internalised bacteria. Many clinically relevant opportunistic pathogens, including *Legionella* [63, 81], *Burkholderia cenocepacia* [82] and *Cryptococcus neoformans* [83], have developed the ability to subvert normal phagosome maturation so as to maintain a permissive niche inside host phagocytes. This ability is likely to have evolved from their ancestors' interactions with environmental predators such as amoebae [84–86].

*Legionella* are phagocytosed in the lung by alveolar macrophages. After internalisation, they employ effectors secreted via their Type IV secretion system, some of which interfere with PI(3)P-signalling, to inhibit phagosome maturation [67, 87, 88]. We have shown that the products of PIKfyve are not required for *Legionella* to establish an intracellular replication niche. Rather, *Legionella* survive much better in PIKfyve-deficient cells, suggesting that PI(3,5)P_2_ helps *Dictyostelium* to eliminate rather than harbour *Legionella*.

This is in contrast to the non-phagocytic invasion of epithelia that occurs during *Salmonella* infection. In that case, PIKfyve activity is necessary to promote the generation of a specialised survival niche within which the bacteria replicate [51]. *Salmonella* has thus evolved a specific requirement for PIKfyve in generating a survival niche – likely through using phagosome acidification as a cue for virulence factor expression – whereas *Legionella* and other bacteria are suppressed by rapid and PIKfyve-driven phagosomal maturation.

The molecular arms race between host and pathogens is complex and of great importance. The very early events of phagosome maturation are critical in this competition; host cells aim to kill their prey swiftly whilst pathogens try to survive long enough to escape. Although its molecular effectors remain unclear, PIKfyve and its products are crucial to tip the balance in favour of the host, providing a general mechanism to ensure efficient antimicrobial activity.

## Materials and Methods

### Cell strains and culture

*Dictyostelium discoideum* cells were grown in adherent culture in plastic Petri dishes in HL5 medium (Formedium) at 22°C. *PIKfyve*^-^ mutants were generated in both Ax2 and Ax3 backgrounds, with appropriate wild-type controls used in each case. Cells were transformed by electroporation and transformants selected in 20 μg/ml hygromycin (Invitrogen) or 10 μg/ml G418 (Sigma). Apilimod was from United States Biological.

Growth in liquid culture was measured by seeding log phase cells in a 6 well plate and counting cells every 12 hours using a haemocytometer. Growth on bacteria was determined by plating ~10 *Dictyostelium* cells on SM agar plates (Formedium) spread with a lawn of non-pathogenic KpGe *K. pneumoniae* [89].

Plaque assays were performed as previously described [90]. Briefly, serial dilution of *Dictyostelium* cells (10–10^4^) were placed on bacterial lawns and grown until visible colonies were obtained. The bacterial strains were kindly provided by Pierre Cosson and were: *K. pneumoniae* laboratory strain and 52145 isogenic mutant (Benghezal et al., 2006), the isogenic *P. aeruginosa* strains PT5 and PT531 (*rhlR-lasR* avirulent mutant) (Cosson et al., 2002), *E.coli* DH5α (Fisher Scientific), *E. coli* B/r (Gerisch, 1959), non-sporulating *B. subtilis* 36.1 (Ratner and Newell, 1978), and *M. luteus* (Wilczynska and Fisher, 1994). An avirulent strain of *K. pneumoniae* was obtained from ATCC (Strain no. 51697).

The *Dictyostelium* development was performed by spreading 10^7^ amoebae on nitrocellulose filters (47 mm Millipore) on top of absorbent discs pre-soaked in KK2 (0.1 M potassium phosphate pH 6.1) and images were taken at 20 hours [91].

### Gene disruption and molecular biology

*PIKfyve*^-^ cells in an Ax2 background were generated by gene disruption using homologous recombination. A blasticidin knockout cassette was made by amplifying a 5' flanking sequence of the *PIKfyve* gene (DDB_G0279149) (primers: fw-GGTAGATGTTTAGGTGGTGAAGT, rv-gatagctctgcctactgaagCGAGTGGTGGAATTCATAAAGG) and 3' flanking sequence (primers: fw-ctactggagtatccaagctgCCATTCAAGATAGACCAACCAATAG, rv-AGAATCAGAATAAACATCACCACC). These primers contained cross over sequences (in lower case) allowing a LoxP-flanked blasticidin resistance cassette (from pDM1079, a kind gift from Douwe Veltman) to be inserted between the two arms.

For *PIKfyve* gene disruption in an Ax3 background a knockout cassette was constructed in pBluescript by sequentially cloning fragment I (amplified by TAGTAGGAGCTCGGATCCGGTAGATGTTTAGGTGGTGAAGTTTTACCAAC and TAGTAGTCTAGACGAGTGGTGGAATTCATAAAGGTACGTTCAT) and fragment II (amplified by TAGTAGAAGCTTCCATTCAAGATAGACCAACCAATAGTAGTCCTGC and TAGTAGGGTACCGGATCCCAGTGTGTAAATGAGAATCAGAATAAACATCACC). The blasticidin resistance gene was inserted between fragment I and II as an Xbal - Hindlll fragment derived from pBSRδBam [92]. Both constructs were linearised, electroporated into cells and colonies were screened by PCR. GFP-2xFYVE was expressed using pJSK489 [41] and GFP-PH_CRAC_ with pDM631 [93]. VatM and VatB were cloned previously [94] but subcloned into the GFP-fusion expression vectors pDM352 and 353 [95] to give plasmids pMJC25 and pMJC31 respectively.

### Microscopy and image analysis

Fluorescence microscopy was performed on a Perkin-Elmer Ultraview VoX spinning disk confocal microscope running on an Olympus 1×81 body with an UplanSApo 60x oil immersion objective (NA 1.4). Images were captured on a Hamamatsu C9100-50 EM-CCD camera using Volocity software by illuminating cells with 488 nm and 594 nm laser lines. Quantification was performed using Image J (https://imagej.nih.gov).

To image PI(3)P dynamics, cells were incubated in HL5 medium at 4 °C for 5 mins before addition of 10 μl of washed 3 μm latex beads (Sigma) and centrifugation at 280 x *g* for 10 seconds in glass-bottomed dishes (Mat-Tek). Dishes were removed from ice and incubated at room temperature for 5 mins before imaging. Images were taken every 30 s across 3 fields of view for up to 30 mins.

V-ATPase recruitment and acidification was performed using *Saccharomyces cerevisiae* labelled with pHrodo red (Life Technologies) as per the manufacturers instructions. *Dictyostelium* cells in HL5 medium were incubated with 1×10^7^ yeast per 3 cm dish, and allowed to settle for 10 mins before the medium was removed and cells were gently compressed under a 1% agarose/HL5 disk. Images were taken every 10 s across 3 fields of view for up to 20 mins. Yeast particles were identified using the "analyse particles" plugin and mean fluorescence measured over time. V-ATPase recruitment was measured as the mean fluorescence within a 0.5 μm wide ring selection around the yeast. The signal was then normalised to the initial fluorescence after yeast internalisation for each cell.

### Endocytosis and exocytosis

To measure endocytosis, *Dictyostelium* at 5 × 10^6^ cells/ml were shaken at 180 rpm for 15 mins in HL5 before addition of 100 mg/ml FITC dextran (molecular mass, 70 kDa; Sigma). At each time point 500 μl of cell suspension were added to 1 ml ice-cold KK2 on ice. Cells were pelleted at 800 x *g* for 2 mins and washed once in KK2. The pellet was lysed in 50 mM Na_2_HPO_4_ pH 9.3 0.2% Triton X-100 and measured in a fluorimeter. To measure exocytosis, cells were prepared as above and incubated in 2 mg/ml FITC-dextran overnight. Cells were pelleted, washed twice in HL5 and resuspended in HL5 at 5 × 10^6^ cells/ml. 500 μl of cell suspension were taken for each time point and treated as described above.

### Phagocytosis and phagosomal activity assays

Phagocytosis of *E. coli* was monitored by the decrease in turbidity of the bacterial suspension over time as described [96]. An equal volume of 2 × 10^7^ *Dictyostelium* cells was added to a bacterial suspension with an OD_600_ nm of 0.8, shaking at 180 rpm, and the decrease in OD_600_ nm was recorded over time. Phagocytosis of GFP-expressing *M. smegmatis* and 1 μm YG-carboxylated polystyrene beads (Polysciences Inc.) was previously described [53, 97]. 2 × 10^6^ *Dictyostelium*/ml were shaken for 2 hours at 150 rpm. Either 1 μm beads (at a ratio of 200:1) or *M. smegmatis* (multiplicity of infection (MOI) 100) were added, 500 μl aliquots of cells were taken at each time point and fluorescence was measured by flow cytometry [53].

Phagosomal pH and proteolytic activity were measured by feeding cells either FITC/TRITC or DQgreen-BSA/Alexa 594-labelled 3 μm silica beads (Kisker Biotech) [53]. Briefly, cells were seeded in a 96 well plate before addition of beads and fluorescence measured on a plate reader over time. pH values were determined by the ratio of FITC to TRITC fluorescence using a calibration curve and relative proteolysis was normalised to Alexa594 fluorescence. To measure proteolytic activity in cell lysates 4 × 10^7^ cells/ml were resuspended in 150 mM potassium acetate pH 4.0 and lysed by 3 cycles of freeze/thaw in liquid nitrogen. After pelleting cell debris at 15000 rpm for 5 minutes at 4°C, 100 μl of lysate was added per well. Proteolytic activity was measured on a plate reader in triplicate, as described above, using 1 × 10^8^ DQ-BSA/Alexa594 beads per cell. A 5 × final concentration of HALT protease inhibitor cocktail (Life Technologies) was added to samples as a negative control.

### Phagosome isolation and blotting

*Dictyostelium* phagosomes were purified at different stages in maturation after engulfment of latex beads as previously described [98]. Briefly 10^9^ cells per timepoint were incubated with a 200-fold excess of 0.8 μm diameter beads (Sigma) first in 5 ml ice-cold Soerensen buffer containing 120 mM sorbitol (SSB) pH 8 for 5 minutes, then added to 100 ml room-temperature HL5 medium in shaking culture (120 rpm) for 5 (first time point) or 15 minutes to allow phagocytosis (pulse). Engulfment was stopped by adding cells to 300ml ice-cold SSB and centrifugation. After washing away non-engulfed beads, cells were again shaken in room-temperature HL5 for the times indicated (chase) to allow maturation. At each time point, maturation was stopped using ice-cold SSB as above, and cells pelleted. Phagosomes were purified as in [56], after homogenization using 10-passages through a ball homogeniser (void clearance 10 μm). The homogenate was incubated with 10 mM Mg-ATP (Sigma) for 15 minutes before loading onto a discontinuous sucrose gradient. Phagosomes were collected from the 10–25% interface, normalised by light scattering at 600 nm and analysed by Western blot. Antibodies used were anti-VatA mAB 221-35-2 (gift from G. Gerisch), anti-vatM mAb N2 [99]; rabbit anti-cathepsin D [100] and anti-Abp1 [101]. All blots were processed in parallel from the same lysates with identical exposure and processing between cell lines.

### Bacteria killing assay

Killing of GFP-expressing *K. pneumoniae* was measured as previously described [60]. Briefly, 10 μl of an overnight culture of bacteria in 280 μl HL5 was placed in a glass-bottomed dish and allowed to settle before careful addition of 1.5 ml of a *Dictyostelium* cell suspension at 1 × 10^6^ cells/ml. Images were taken every 20 s for 40 min at 20x magnification. Survival time was determined by how long the GFP-fluorescence persisted after phagocytosis.

### Western blotting

Ax2 or *PIKfyve*^-^ cells expressing GFP-VatM or VatB-GFP were analysed by SDS-PAGE and Western blot using a rabbit anti-GFP primary antibody (gift from A. Peden) and a fluorescently conjugated anti-rabbit 800 secondary antibody, using standard techniques. The endogenous biotinylated mitochondrial protein Methylcrotonoyl-CoA Carboxylase 1 (MCCC1) was used as a loading control using Alexa680-conjugated Streptavadin (Life Technologies) [102].

### *Legionella* infection assays

The following *L. pneumophila* Philadelphia-1 strains were used: virulent JR32 [103], the isogenic *∆icmT* deletion mutant GS3011 lacking a functional Icm/Dot type IV secretion system [73], and corresponding strains constitutively producing GFP [74]. *L. pneumophila* was grown for 3 days on charcoal yeast extract (CYE) agar plates, buffered with *N*-(2-acetamido)-2-aminoethane-sulfonic acid (ACES) [104]. For infections, liquid cultures were inoculated in AYE medium at an OD_600_ of 0.1 and grown for 21 h at 37 °C (post-exponential growth phase). To maintain plasmids, chloramphenicol was added at 5 μg/ml.

Uptake by *D. discoideum*, intracellular replication or killing of GFP-producing *L. pneumophila* was analyzed by flow cytometry as described [71]. Exponentially growing amoebae were seeded onto a 24-well plate (1 × 10^6^ cells/ml HL5 medium per well) and allowed to adhere for at least 1–2 h. *L. pneumophila* grown for 21 h in AYE medium was diluted in HL5 medium and used to infect the amoebae at an MOI of 50. The infection was synchronized by centrifugation (10 min, 500 x *g*), and infected cells were incubated at 25 °C for 30 min before extracellular bacteria were removed by washing twice with SorC (2 mM Na_2_HPO_4_,15 mM KH_2_PO_4_, 50 μM CaCl_2_, pH 6.0). Infected amoebae were detached by vigorously pipetting and fixed (PFA 2%, sucrose 125 mM, picric acid 15%, in PIPES buffer, pH 6.0), and 1 × 10^4^ amoebae per sample were analyzed using a using a LSR II Fortessa analyser. The GFP fluorescence intensity falling into a *Dictyostelium* scatter gate was quantified using FlowJo software (Treestar, http://www.treestar.com).

Alternatively, intracellular replication of *L. pneumophila* in *D. discoideum* was quantified by determining colony forming units (CFU) in the supernatant as described [71, 105]. Exponentially growing amoebae were washed and resuspended in MB medium (7 g of yeast extract, 14 g of thiotone E peptone, 20 mM MES in 11 of H_2_O, pH 6.9). Amoebae (1 × 10^5^ cells per well) were seeded onto a 96-well plate, allowed to adhere for at least 2 h, and infected at an MOI of 0.1 with *L. pneumophila* grown in AYE medium for 21 h and diluted in MB medium. The infection was synchronized by centrifugation, and the infected amoebae were incubated at 25°C. At the time points indicated, the number of bacteria released into the supernatant was quantified by plating aliquots (10–20 μl) of appropriate dilutions on CYE plates. Intracellular bacteria were also quantified by counting CFU after lysis of the infected amoebae with saponin. At the time points indicated host cells were detached by vigorous pipetting and lysed by incubation with saponin (final concentration - 0.8%) for 10 min. The number of bacteria released into the supernatant was quantified by plating 20 μl aliquots of appropriate dilutions on CYE plates.

## Acknowledgments

The authors would like to Phill Hawkins for his endeavours to separate *Dictyostelium* PIP_2_s

---

**Supplementary Figure 1.**
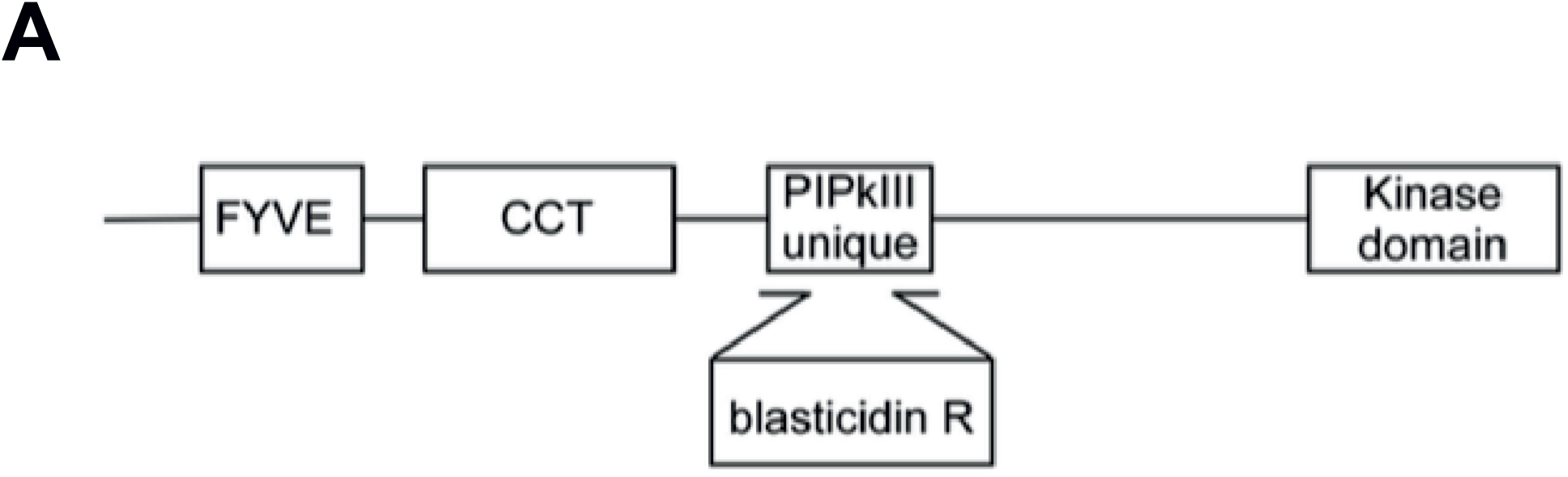
*PIKfyve* gene disruption. (A) Schematic representation of *Dictyostelium* PIKfyve indicating the site where the blasticidin resistance cassette was inserted into the gene to generate knockouts.

**Supplementary Figure 2.**
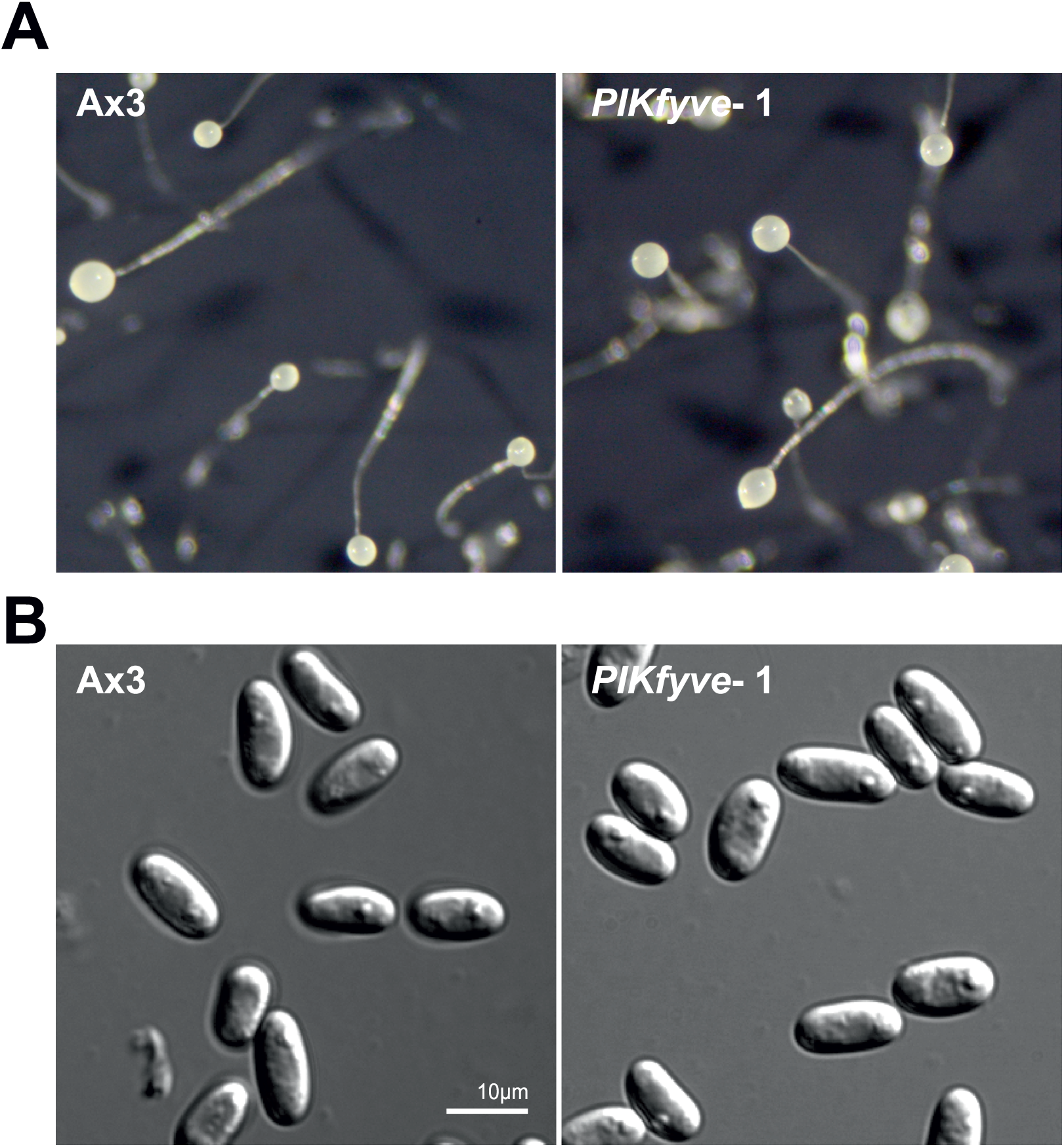
PIKfyve is not required for development. (A) Images of *Dictyostelium* fruiting bodies formed on filter discs, indicating a normal morphology and proportioning in the absence of PIKfyve. (B) Higher magnification differential interference contrast (DIC) images of pores collected from the fruiting bodies in (A).

**Supplementary Figure 3.**
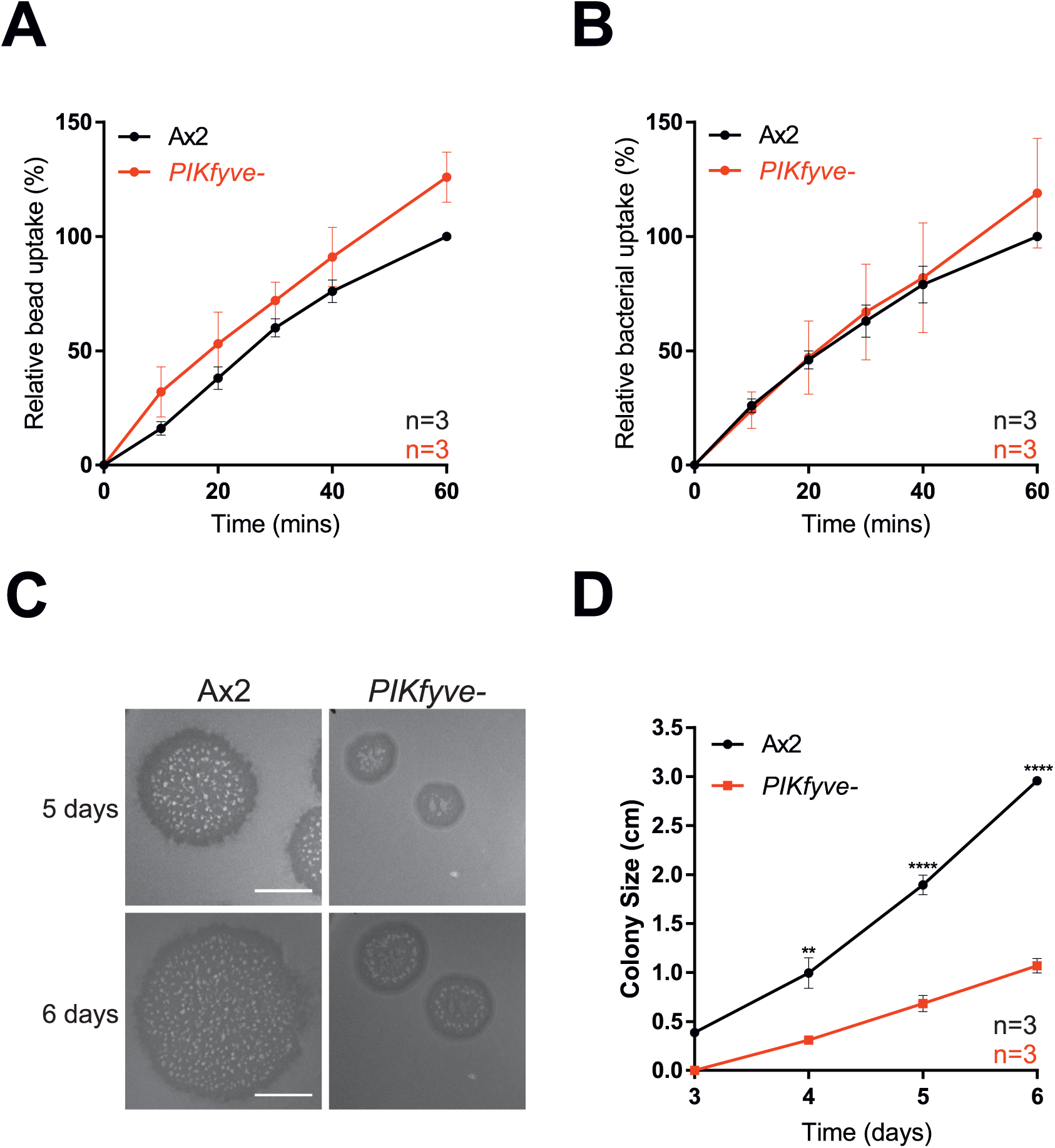
Conservation of *PIKfyve*-null phenotypes in Ax2-derived mutants. Phagocytosis of (A) of 1 μm beads or (B) GFP-expressing *Mycobacterium smegmatis* measured by flow cytometry, is normal in *PIKfyve^-^*null cells. (C) Growth on lawns of *K. pneumoniae* is impaired. Colony diameter over time is plotted in (D). All data are means +/-SD.

**Supplementary Figure 4.**
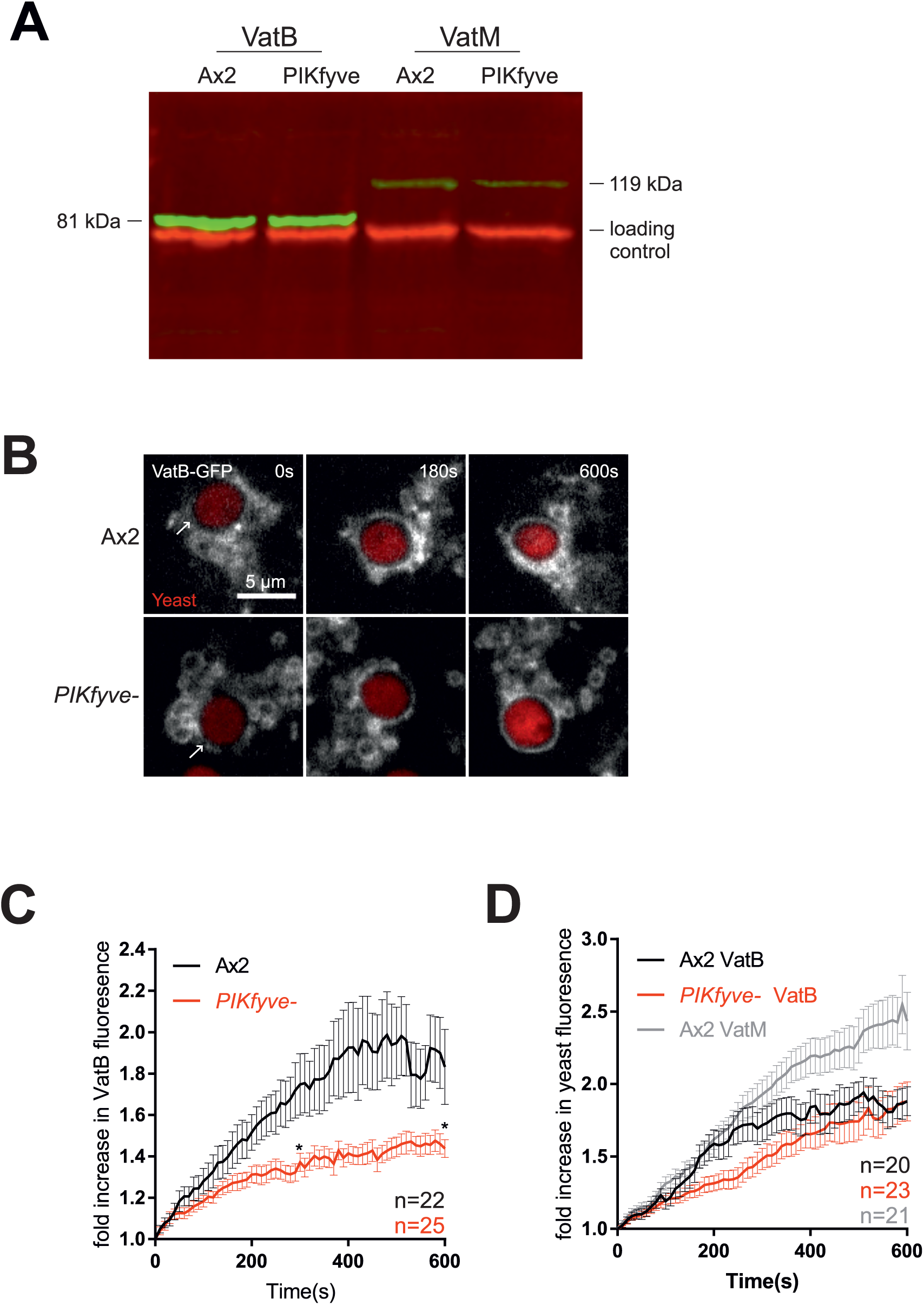
VatB-GFP expression has a dominant negative effect on acidification. (A) Western blot of cells expressing VatB-GFP or GFP-VatM, probed with an anti-GFP antibody (green). There was no difference in expression levels between Ax2 and *PIKfyve*^-^ cells for either reporter. However, VatB-GFP was expressed at higher levels than GFP-VatM, likely because it is present in 3 copies per V-ATPase complex. Loading control is the mitochondrial protein MCCC1, recognised by Alexa680-conjugated streptavidin (red). (B) Recruitment of vatB-GFP to phagosomes containing pHrodo-labelled yeast. (C) Automated image analysis of vatB-GFP recruitment as described in Figure 3, showing reduced recruitment in *PIKfyve^-^*null cells. (D) Phagosome acidification, measured by the increase in pHrodo fluorescence over time. Note that expression of vatB-GFP in Ax2 cells significantly reduces phagosome acidification relative to GFP-vatM expressing cells, indicating disruption of V-ATPase activity. Values plotted are mean +/- SEM.

